# Engineered basement membrane mimetic hydrogels to study mammary epithelial morphogenesis and invasion

**DOI:** 10.1101/2025.02.28.640825

**Authors:** Jane A. Baude, Megan D. Li, Sabrina M. Jackson, Abhishek Sharma, Daniella I. Walter, Ryan S. Stowers

## Abstract

Reconstituted basement membrane (rBM) products like Matrigel are widely used in 3D culture models of epithelial tissues and cancer. However, their utility is hindered by key limitations, including batch variability, xenogenic contaminants, and a lack of tunability. To address these challenges, we engineered a 3D basement membrane (eBM) matrix by conjugating defined extracellular matrix (ECM) adhesion peptides (IKVAV, YIGSR, RGD) to an alginate hydrogel network with precisely tunable stiffness and viscoelasticity. We optimized the mechanical and biochemical properties of the engineered basement membranes (eBMs) to support mammary acinar morphogenesis in MCF10A cells, similar to rBM. We found that IKVAV-modified, fast-relaxing (τ_1/2_ = 30-150 s), and soft (E = 200 Pa) eBMs best promoted polarized acinar structures. Clusters became invasive and lost polarity only when the IKVAV-modified eBM exhibited both similar stiffness to a malignant breast tumor (E = 4000 Pa) and slow stress relaxation (τ_1/2_ = 600-1100 s). Notably, tumor-like stiffness alone was not sufficient to drive invasion in fast stress relaxing matrices modified with IKVAV. In contrast, RGD-modified matrices promoted a malignant phenotype regardless of mechanical properties. We also utilized this system to interrogate the mechanism driving acinar and tumorigenic phenotypes in response to microenvironmental parameters. A balance in activity between β1- and β4-integrins was observed in the context of IKVAV-modified eBMs, prompting further investigation into the downstream mechanisms. We found differences in hemidesmosome formation and production of endogenous laminin in response to peptide type, stress relaxation, and stiffness. We also saw that inhibiting either focal adhesion kinase or hemidesmosome signaling in IKVAV eBMs prevented acinus formation. This eBM matrix is a powerful, reductionist, xenogenic-free system, offering a robust platform for both fundamental research and translational applications in tissue engineering and disease modeling.

## Introduction

The three-dimensional (3D) microenvironment surrounding mammary epithelium profoundly influences both the normal morphogenesis of these cells and the progression of breast tumors (1–5). Throughout development and into adulthood, the mammary epithelium undergoes continuous and dynamic morphogenetic changes, often accompanied by significant remodeling of the extracellular matrix (ECM).

Alterations in ECM composition and mechanical properties play a critical role in modulating these morphogenetic processes, including epithelial-to-mesenchymal transition, and are key drivers of tumor progression (6–8). However, due to the ECM’s complexity and its dynamic, interactive components, fully understanding the interplay between ECM behavior and morphogenetic programs requires the development of a controllable and tunable in vitro platform.

In vivo, basement membrane proteins of the ECM directly contact mammary epithelium cells (MECs) and govern their phenotype and polarization. The majority of in vitro models of mammary morphogenesis rely on reconstituted basement membrane (rBM) extract (i.e. Matrigel™, Cultrex™, Geltrex™) to recapitulate the epithelial basement membrane (9). Researchers first established the use of 3D rBM culture for MECs in studies that emphasized the important role of rBM in facilitating mammary morphogenesis in vitro (1,10). As a result, rBM has become a cornerstone for investigating breast cancer mechanobiology, with landmark studies (11–18) demonstrating how the mechanical properties of the extracellular matrix can drive tumor progression. The tumor microenvironment is significantly stiffer than healthy tissue, and studies have shown that matrix stiffness is a key mechanical property that enhances the tumorigenic behavior of mammary epithelial cells in rBM systems (18,19). 3D in vitro studies have shown that kilopascal stiffness induces malignant phenotypes in MECs while soft substrates (< 200 Pa) support a non-malignant mammary acinar phenotype (17). While the literature extensively documents the impacts of matrix stiffness in vivo and in vitro (17,18,20–22), viscoelasticity has received less attention in the context of mammary epithelium, though recent studies demonstrate that viscoelasticity can impact mammary epithelial and breast cancer cell phenotypes (23–25). Although rBM is inherently viscoelastic, the field has only recently started to investigate these properties within 3D culture systems (26–30).

Despite its widespread use, rBM products suffer from several limitations, including batch-to-batch variability, a lack of mechanical and biochemical tunability, the presence of undefined growth factors and signaling molecules, xenogenic contaminants, and, recently, pandemic-related supply shortages (31–33). These limitations have driven the development of engineered matrices that mimic key features of rBM. Hydrogels, particularly those derived from bioinert materials, offer a versatile platform. While engineered rBM-free systems have shown promise in other organoid models, such as intestinal and neuroepithelial (34–37), fully recapitulating mammary morphogenesis without rBM remains a substantial challenge. Deciphering the mechanisms by which biochemical and mechanical cues, both independently and in concert, can influence mammary epithelial cell behavior is crucial to achieve this goal. This underscores a two-fold critical need; first, to develop an rBM-free system that effectively supports mammary morphogenesis, and second, to ensure that the system is defined and tunable, which allows for a systematic exploration of the interplay between mechanical and biochemical cues.

In this study, we endeavored to develop a defined, engineered rBM-free matrix system to promote mammary epithelium cell acinar morphogenesis and invasion in vitro. To achieve this, we conjugated a panel of basement membrane-derived adhesion motifs to a mechanically tunable alginate hydrogel network. This innovative, bottom-up approach allows us to precisely and independently manipulate both the biochemical and mechanical attributes of the matrix. By encapsulating MECs within these matrices, we gain a clearer understanding of how the distinct biochemical and mechanical properties influence mammary epithelium cell phenotype individually. Remarkably, our platform offers critical insights into mammary epithelial cell adhesion and morphogenesis, mechanotransduction, and basement membrane production that traditional rBM systems often overlook, illuminating the complex interplay of cellular behavior within a finely controlled environment.

## Results

### Alginate Networks Can be Mechanically and Biochemically Tuned

Using an alginate-based system, matrix stiffness, stress relaxation rate, adhesion ligand type and density can be tuned independently of each other, without altering matrix pore size and architecture (38) (Fig. 1A). Alginate presents no cell adhesion motifs recognized by mammalian cells, thus cell adhesion cues can be entirely controlled by conjugating synthetic adhesion peptide sequences to the alginate chains. Laminin 111 is one of the most abundant and critical proteins found in the mammary basement membrane and in rBM products (39–44). Levels of laminin 111 fluctuate significantly during mammary morphogenesis and the progression of breast cancer (45,46), suggesting that laminin 111 adhesion motifs are promising candidates for modeling the basement membrane. Thus, we selected the laminin 111 adhesion motifs IKVAV and YIGSR to minimally mimic key motifs of the basement membrane and rBM (47). We also opted to use an RGD motif as it is abundant amongst fibronectin and laminin 111 molecules within the basement membrane proteins and frequently used in engineered hydrogel platforms for cell adhesion (14,48–54). We used carbodiimide chemistry (48) to couple peptide sequences containing the adhesion motifs IKVAV, YIGSR, or RGD to purified alginate, at both high and low concentrations (400 and 50 μM, respectively), to test the impacts of changing ligand density (Fig. 1A, B). We verified peptide conjugation via H-NMR (Fig. S1, Table S1).

**Fig. 1.**
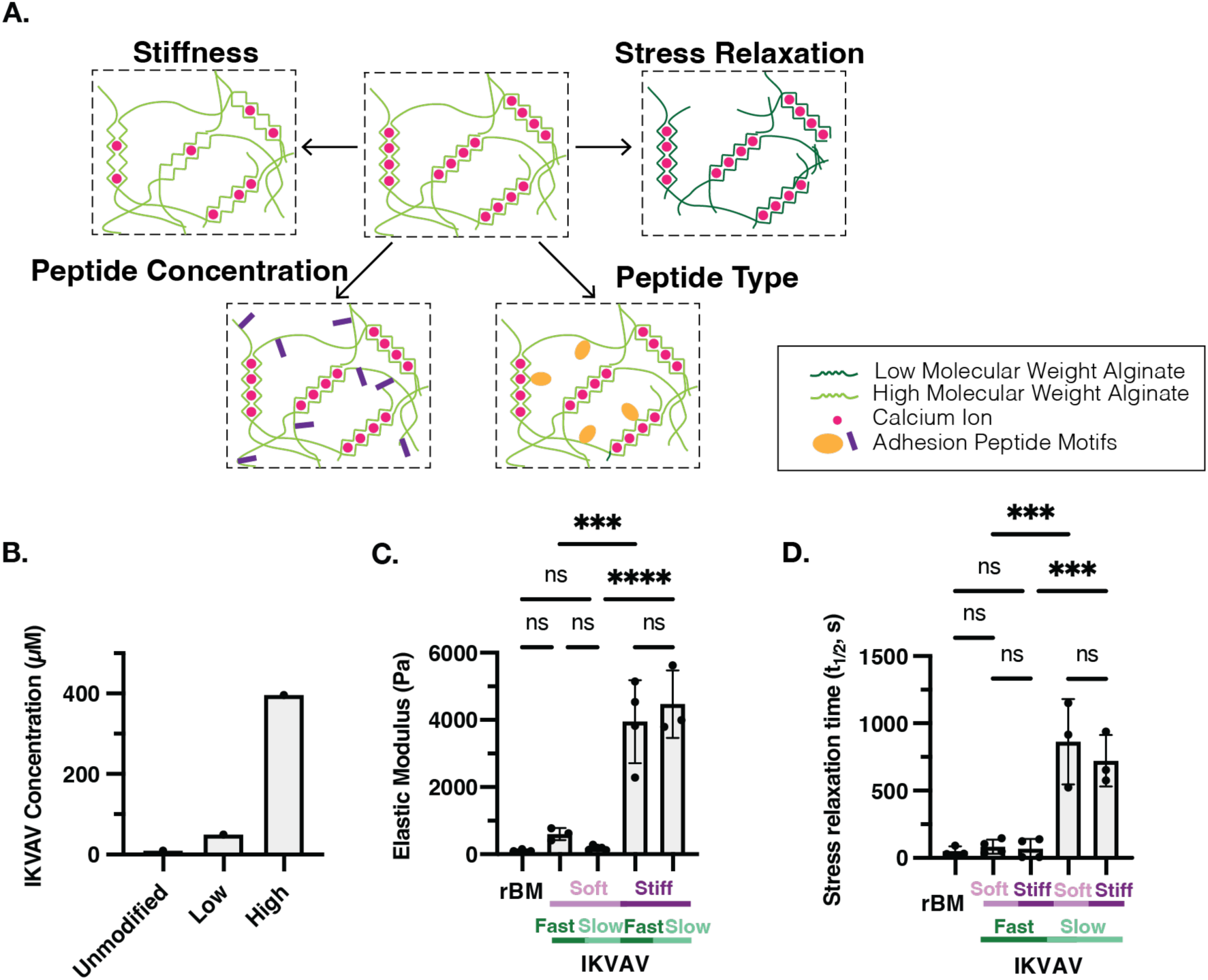
Modified eBMs Have Independently Tunable Mechanical and Biochemical Properties. (**A**) Alginate-based eBMs have tunable mechanical and biochemical properties. Stiffness can be modulated by adjusting the concentration of calcium for crosslinking. Stress relaxation can be changed by utilizing alginates of different molecular weights. Peptides of different types or concentrations can be conjugated to the alginate network. (**B**) Summary of concentration of IKVAV peptide on modified fastrelaxing alginate from H-NMR data (Fig. S1, Table S1). (**C**) Elastic modulus of rBM and IKVAV-modified eBMs. (**D**) Stress relaxation rate of rBM and IKVAV-modified eBMs. Significance was determined by oneway ANOVA for elastic modulus and stress relaxation time (p-value < 0.05). n=1 replicate per condition for H-NMR measurements. n = 3-4 replicates for elastic modulus and stress relaxation time. * indicates p < 0.05, ** p < 0.01, *** p < 0.001, **** p < 0.0001, ns= not significant.

Matrix stiffness, or the resistance to deformation, is known to impact MCF10A phenotype in 3D cell culture studies (14,55). Therefore, it is crucial to have independent control of matrix stiffness relative to other mechanical properties like stress relaxation or pore size within this system. The crosslinking density, and thus stiffness, of alginate hydrogels is a function of the concentration of divalent cations, such as Ca^2+^, used to generate ionic crosslinks. We optimized calcium concentrations for peptide-modified alginate matrices to mimic the reported elastic moduli of either normal mammary gland tissue (< 500 Pa, “soft”), or primary breast tumors (4,000 Pa, “stiff”) (Fig. 1C).

We measured the stress relaxation half-time (τ_1/2_) of rBM to be approximately 20-90 s, indicating it is highly viscoelastic (Fig. 1D). To vary stress relaxation rates in alginate matrices, we used alginates of different molecular weights (38). Low molecular weight alginate modified with IKVAV yielded similar stress relaxation profiles as rBM (τ_1/2_ = 30-150 s) (Fig. 1D). In contrast, high molecular weight alginate modified with IKVAV produced slow-relaxing matrices (τ_1/2_ = 600-1100 s) (Fig. 1D) (56). Importantly, the elastic modulus and stress relaxation rate can be independently tuned (Fig. 1C, D). Statistical analysis showed no significant differences in stiffness between matrices with different stress relaxation rates, nor in stress relaxation rates between matrices of different stiffness; both soft and stiff matrices to be created with either fast or slow stress relaxation (Fig. 1C, D).

### Soft, Fast-Relaxing Modified Matrices Enable Mammary Acinus Morphogenesis Without rBM

Mammary acini are three-dimensional structures that develop within a seven to fourteen-day period, distinguished by their apical-basal polarity and hollow lumens (1).

MCF10A breast epithelial cells encapsulated in rBM develop into polarized acini, mimicking in vivo mammary acini (10). To assess whether acinar morphogenesis of MCF10A cells could occur in modified eBMs, we encapsulated MCF10A cells matrices that most closely mimicked the properties of rBM, namely a high concentration of adhesion peptides, fast-relaxation, and soft modulus. Within these soft, fast-relaxing matrices, we varied the peptide motifs to create three distinct conditions: (1) IKVAV-modified, (2) YIGSR-modified, and (3) RGD-modified eBMs.

The cells were cultured in the respective matrix condition for two weeks and then cell cluster morphology was analyzed and compared to cells cultured in rBM. In IKVAV-modified soft eBMs, MCF10As developed into acinar-like clusters over the two-week culture period, very similar to cells grown in rBM (Fig. 2A). MCF10A cells cultured in YIGSR-modified eBMs also formed round clusters that closely resembled those observed in rBM, though with less frequency than in IKVAV-modified matrices (Fig. 2A). In contrast, MCF10As encapsulated in RGD-modified soft matrices remained as single cells or small, irregularly shaped clusters (Fig. 2A). In order to further validate the extent of acinus formation and maturation in these matrices we performed immunostaining and confocal microscopy for key markers of polarization; β4-integrin (basal) and Golgi matrix protein 130 (GM130, apical). In IKVAV-modified fast-relaxing, soft eBMs we observed similar polarization to that of cell clusters in rBM (Fig. 2B). For instance, β4-integrin localized to the basement membrane in both the IKVAV-modified matrix and rBM, while GM130 was apical to the nuclei of the cluster in both conditions (Fig. 2B). Of note, in YIGSR-modified fast-relaxing, soft eBMs polarization was less pronounced than IKVAV-modified matrices, despite the rounded clusters (Fig. 2B, C). In the RGD-modified matrices both β4-integrin and GM130 were distributed throughout the cluster, indicating a lack of polarization (Fig. 2B).

**Fig. 2.**
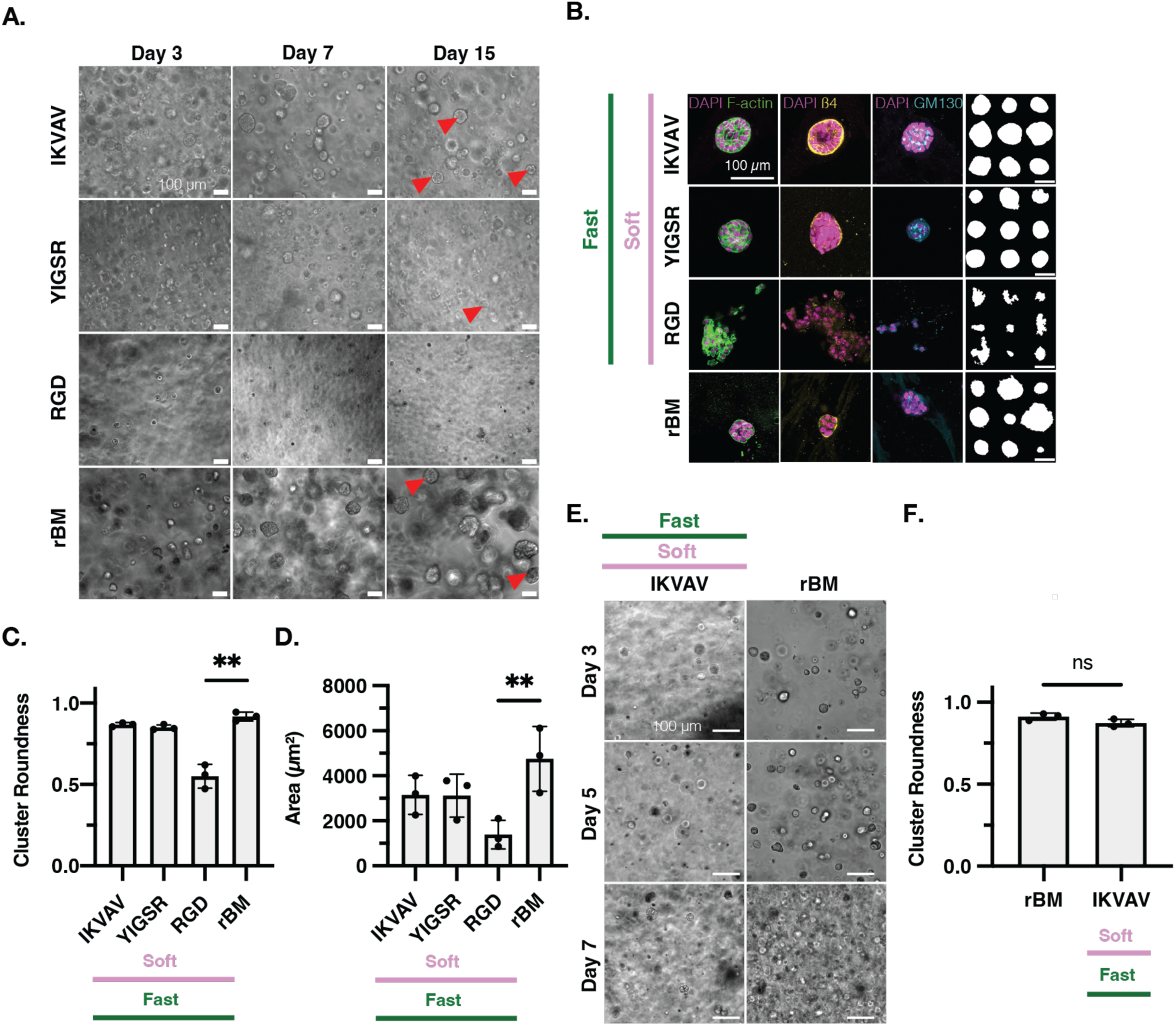
Modified-alginate eBMs mimicking rBM enable mammary acinus formation and organoid morphogenesis without rBM. (**A**) Representative brightfield images of MCF10As in fast-relaxing and soft eBMs, modified with either IKVAV, YIGSR, or RGD, in comparison to MCF10As grown in rBM. Images are from Day 3, Day 7, and Day 15 during culture. Red arrowheads point to acinar-like clusters. (**B**) Representative confocal images of MCF10As in fast-relaxing and soft eBMs, modified with either IKVAV, YIGSR, or RGD, compared to MCF10As encapsulated in rBM. From left to right DAPI/F-actin, DAPI/β4-integrin, DAPI/GM130, and representative outlines of the respective condition. Scale bars = 100 µm. (**C**) Quantification of cluster roundness from the fast-relaxing and soft eBMs. (**D**) Representative brightfield images of PDM-350 organoids encapsulated in IKVAV-modified, fast-relaxing, soft eBM compared to rBM. (**E**) Representative brightfield images of PDM-350 organoids in IKVAV-modified eBMs compared to rBM. (**F**) Quantification of cluster roundness from Panel F. All scale bars = 100 µm. Data are shown as mean ± SD of three biological replicates (n=3, the average of 20-50 images per replicate) unless otherwise indicated. Significance was determined by a Kruskal-Wallis test followed by Dunnett’s multiple testing correction for MCF10A cluster roundness and by one-way ANOVA followed by post hoc multiple comparison tests for cluster area. An unpaired t-test was used for PDM-350 cluster roundness. If no statistical significance indicator bars are shown, there were no significant differences (p-value > 0.05). * indicates p < 0.05, ** p < 0.01, *** p < 0.001, **** p < 0.0001, ns= not significant.

We also analyzed overall cell cluster roundness and area. MCF10A clusters in IKVAV- and YIGSR-modified soft eBMs had high roundness values, which were not significantly different from each other, or from those measured of clusters in rBM (Fig. 2C). In contrast, clusters in the RGD-modified, soft eBMs were significantly less round (Fig. 2C). We also compared the sizes of the different clusters. Clusters grown in IKVAV- or YIGSR-modified matrices had similar area to those grown in rBM, while clusters in RGD-modified matrices were significantly smaller (Fig. 2D).

In order to assess the utility of this system as an alternative to rBM for other cell culture studies, we next attempted to grow patient-derived organoids (PDOs) in modified matrices. We chose IKVAV-modified, fast-relaxing, soft matrices as a candidate condition to test the PDO samples since this condition generated the most robust acini formation with MCF10As. Remarkably, PDM-350 derived clusters grown in IKVAV-modified, fast-relaxing, and soft eBMs were very round (Fig. 2E), and not significantly different in cluster roundness from those grown in rBM matrices (Fig. 2F). These results suggest that IKVAV-modified, fast-relaxing, soft eBMs provide a suitable environment for patient-derived organoids, supporting their growth and morphology similarly to traditional rBM matrices.

### Tumor-like Matrix Stiffness Alone Does Not Induce an Invasive Phenotype

After demonstrating that modified matrices could support mammary acinar morphogenesis without rBM in soft, fast-relaxing, IKVAV-modified matrices, we sought to utilize the mechanical tunability of this platform to investigate how changes in stiffness or stress relaxation would impact the phenotype of MCF10A cells. Matrix stiffness in the range of 1-5 kPa is known to induce a tumorigenic phenotype in MCF10As, and we thus hypothesized that modified eBMs with elevated stiffness would drive invasion (55,57). Surprisingly, in the fast-relaxing, IKVAV-modified matrices, we found no significant differences in the cluster area, roundness, or invasiveness of MCF10A cells between the soft and stiff matrices (Fig. 3A-D). In both conditions, acinar development was evident, characterized by clear apical-basal polarity and high roundness (Fig. 3A,B).

**Fig. 3.**
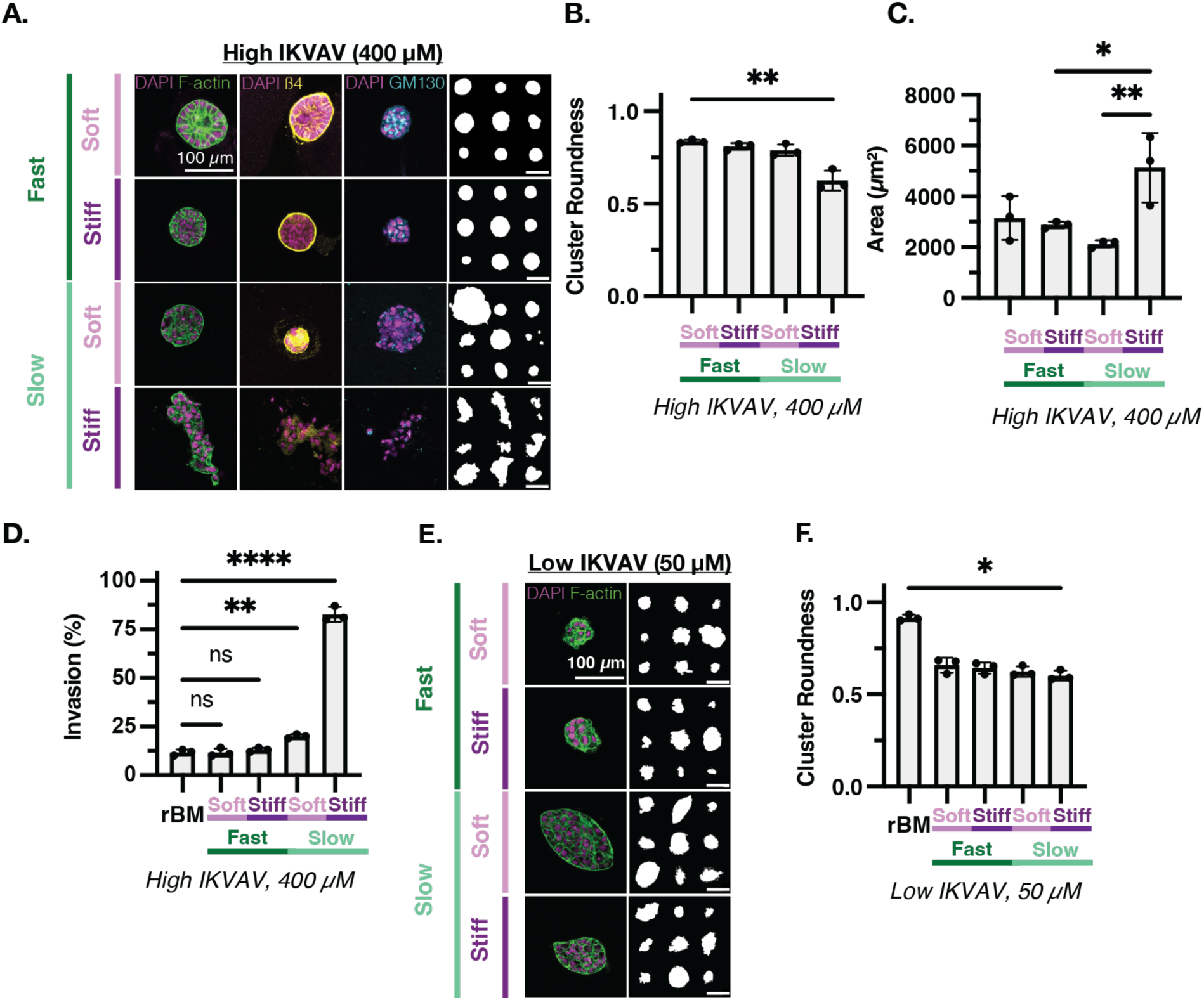
Both elevated stiffness and slow stress relaxation are required to induce invasion in IKVAV-modified eBMs. (**A**) Representative confocal images of MCF10As encapsulated in high concentration IKVAV-modified fast-relaxing and soft, fast-relaxing and stiff, slow-relaxing and soft, and slow-relaxing and stiff eBMs. From left to right: DAPI/F-actin, DAPI/β4-integrin, DAPI/GM130, and representative outlines of the respective condition. (**B**) Quantification of cluster roundness for high concentration IKVAV-modified eBMs. (**C**) Quantification of cluster area for IKVAV-modified eBMs. (**D**) Quantification of the percentage of invasive clusters in IKVAV-modified eBMs. (**E**) Representative confocal images of MCF10As encapsulated in low concentration IKVAV-modified fast-relaxing and soft, fast-relaxing and stiff, slow-relaxing and soft, and slow-relaxing and stiff eBMs. From left to right: DAPI/F-actin and representative outlines. (**F**) Quantification of cluster roundness for low concentration IKVAV-modified eBMs. All scale bars = 100 µm. Data are shown as mean ± SD of three biological replicates (n = 3, 20-50 images per replicate) unless otherwise indicated. Statistical significance was tested by a Kruskal-Wallis test followed by Dunnett’s multiple testing correction for cluster roundness and percentage of invasive clusters, and by one-way ANOVA and post hoc multiple comparison tests for cluster area. If no statistical significance indicator bars are shown, there were no significant differences (p-value > 0.05). * indicates p < 0.05, ** p < 0.01, *** p < 0.001, **** p < 0.0001, ns= not significant.

Although it is established that different tissues have distinct stress relaxation properties (56), and specifically that breast cancer tissue relaxes differently from normal mammary tissue (24), the precise impact of stress relaxation on mammary acinar morphogenesis remains unclear. Therefore, we encapsulated MCF10As in slow-relaxing, IKVAV-modified matrices to compare to IKVAV-modified fast-relaxing eBMs. In slow-relaxing, soft, IKVAV-modified matrices we observed round clusters (Fig. 3A, B), yet these clusters were marked by increased β4-integrin within the cluster as opposed to localizing to the basement membrane, indicating a loss of polarization (Fig. 3A).

Strikingly, when the IKVAV-modified eBMs were both stiff and slow-relaxing, we observed the emergence of multiple invasive regions, loss of polarity and roundness, and increased cluster size (Fig. 3A, D), consistent with prior reports of elevated stiffness driving MCF10A invasion. Furthermore, IKVAV-modified stiff, slow-relaxing matrices had significantly more invasive clusters in comparison to the three other IKVAV conditions tested (Fig. 3D). Thus, in IKVAV eBMs, both a stiff and a slow-relaxing matrix is essential to promote the invasive phenotype. Notably, we observed no noticeable formation of mammary acinar structures in eBMs modified with a low concentration of IKVAV (50 μM) across the mechanical conditions tested. While these clusters were not invasive, they exhibited lower cluster roundness than cells cultured in rBM and were generally more disorganized compared to cells in either rBM or high concentration IKVAV matrices (Fig. 3E, F).

MCF10A cells encapsulated in YIGSR-modified, fast-relaxing, soft eBMs developed acinar-like clusters, but these clusters were less uniform in size and shape and showed less defined lumen formation (Fig. S2A-C), compared to those formed in IKVAV-modified, fast-relaxing, soft eBMs. We also encapsulated MCF10As in YIGSR eBMs that were fast-relaxing and stiff, as well as slow-relaxing and soft, and slow-relaxing and stiff (Fig. S2A-C). We did not see consistent acinar or invasive phenotypes and therefore did not investigate these eBMs further. Additionally, we did not observe any significant mammary morphogenesis in low YIGSR-modified eBMs across the mechanical conditions tested (Fig. S2D,E).

### RGD-modified Matrices Induces Invasiveness, Independent of Mechanical Properties

We next sought to understand how MCF10As behave in RGD-modified matrices.

Notably, MCF10As grown in RGD-modified eBMs did not form acini in any conditions, underscoring the necessity of appropriate adhesion ligands for acinar development. We saw invasive phenotypes in all combinations of mechanical properties (Fig. 4A). Neither β4-integrin nor GM130 localized to the basal or apical membrane, respectively, in MCF10A clusters in RGD eBMs (Fig. 4A), indicating a lack of polarization. MCF10A cluster roundness nor area was significantly different between mechanical conditions of RGD-modified eBMs tested (Fig. 4B, C). Furthermore, the percentage of invasive clusters in RGD eBMs was significantly higher than those found in rBM, with the RGD-modified, slow-relaxing, stiff group being the most invasive (Fig. 4D). After revealing these striking differences in phenotype between IKVAV and RGD eBMs, we were compelled to investigate the molecular mechanisms underlying the variation in MCF10A phenotypes across these modified eBMs.

**Fig. 4.**
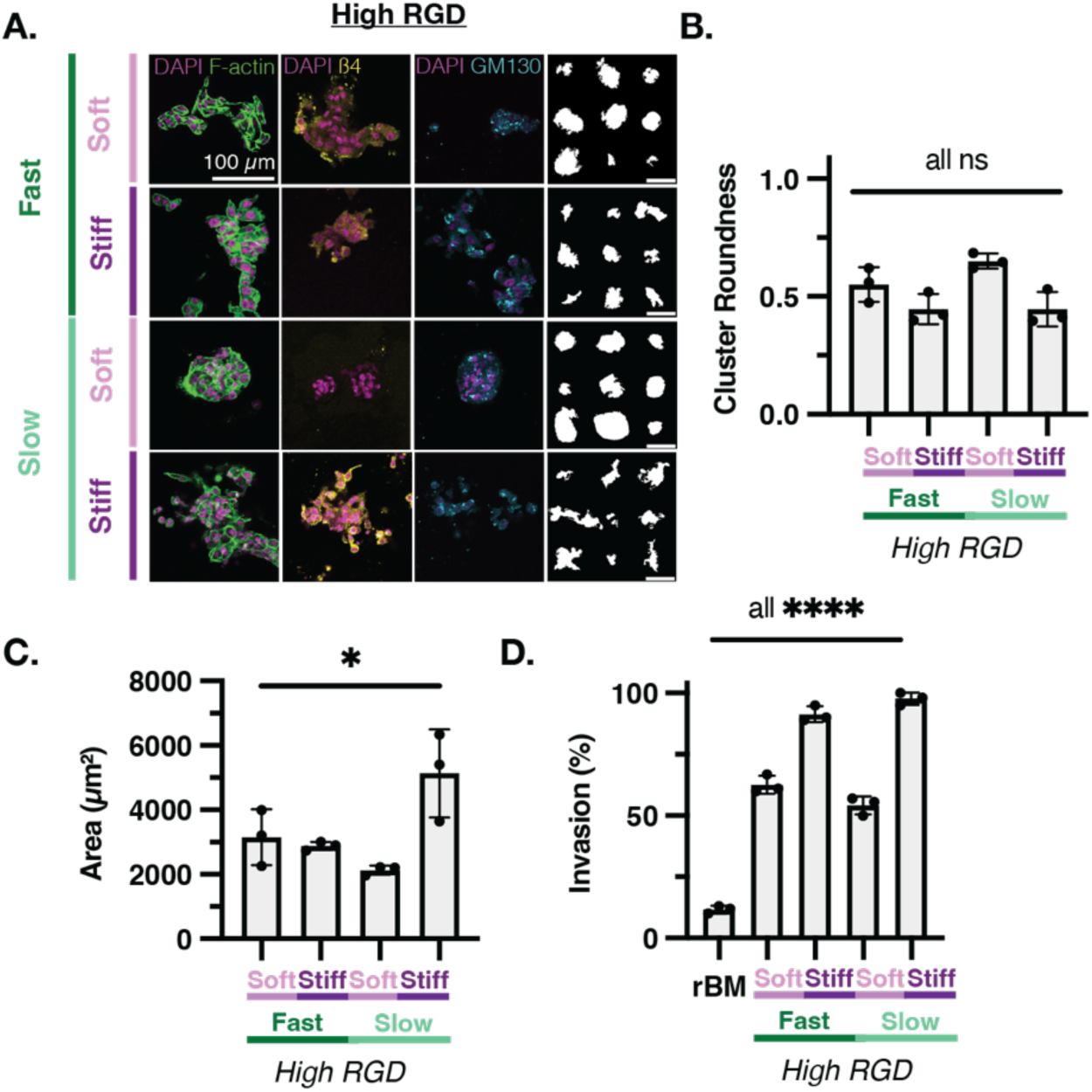
The invasive phenotype is prominent in RGD eBMs regardless of mechanics. (**A**) Representative confocal images of MCF10As encapsulated in high concentration RGD-modified fast-relaxing and soft, fast-relaxing and stiff, slow-relaxing and soft, and slow-relaxing and stiff eBMs. From left to right: DAPI/F-actin, DAPI/β4-integrin, DAPI/GM130, and representative outlines of the respective condition. (**B**) Quantification of cluster roundness for high concentration RGD-modified eBMs. (**C**) Quantification of cluster area for RGD-modified eBMs. (**D**) Quantification of the percentage of invasive clusters in RGD-modified eBMs. All scale bars = 100 µm. Data are shown as mean ± SD of three biological replicates (n=3, 20-50 images per replicate) unless otherwise indicated. Statistical significance was determined by a Kruskal-Wallis test followed by Dunnett’s multiple testing correction for cluster roundness and the percentage of invasive clusters, and by one-way ANOVA and post hoc multiple comparison tests for cluster area. If no statistical significance indicator bars are shown, there were no significant differences (p-value > 0.05). * indicates p < 0.05, ** p < 0.01, *** p < 0.001, **** p < 0.0001, ns= not significant.

### Differential β4- and β1-Integrin Localization Correlates with IKVAV and RGD Peptides

We hypothesized that ligand-specific integrin signaling underlies the divergent phenotypes observed between IKVAV- and RGD-modified eBMs, as these motifs are bound by distinct integrin subunit combinations. IKVAV motifs are predominantly bound by β1-integrins (e.g., α3β1, α2β1, α6β1), while RGD motifs are engaged by αv- and β1-integrins in MECs (58). It is important to note that while β1-integrins are crucial for IKVAV and RGD binding, other integrin combinations also play important roles in MECs. For example, α6β4 integrin is a key receptor for laminin, a major component of the basement membrane, and is essential for hemidesmosome formation and cell adhesion (59).

In IKVAV-modified eBMs, β1- and β4-integrins were basally localized in fast-relaxing matrices (both soft and stiff) as well as in slow-relaxing, soft matrices (Fig. 5A). However, in slow-relaxing, stiff, IKVAV eBMs, β1- and β4-integrins were more randomly distributed with higher variability in signal intensity (Fig. 5A). Unsurprisingly, we also saw dispersed organization of both β1- and β4-integrin in all RGD conditions (Fig. 5A). Furthermore, β1-integrins were significantly more abundant than β4-integrins in RGD eBMs (Fig. S5B).

**Fig. 5.**
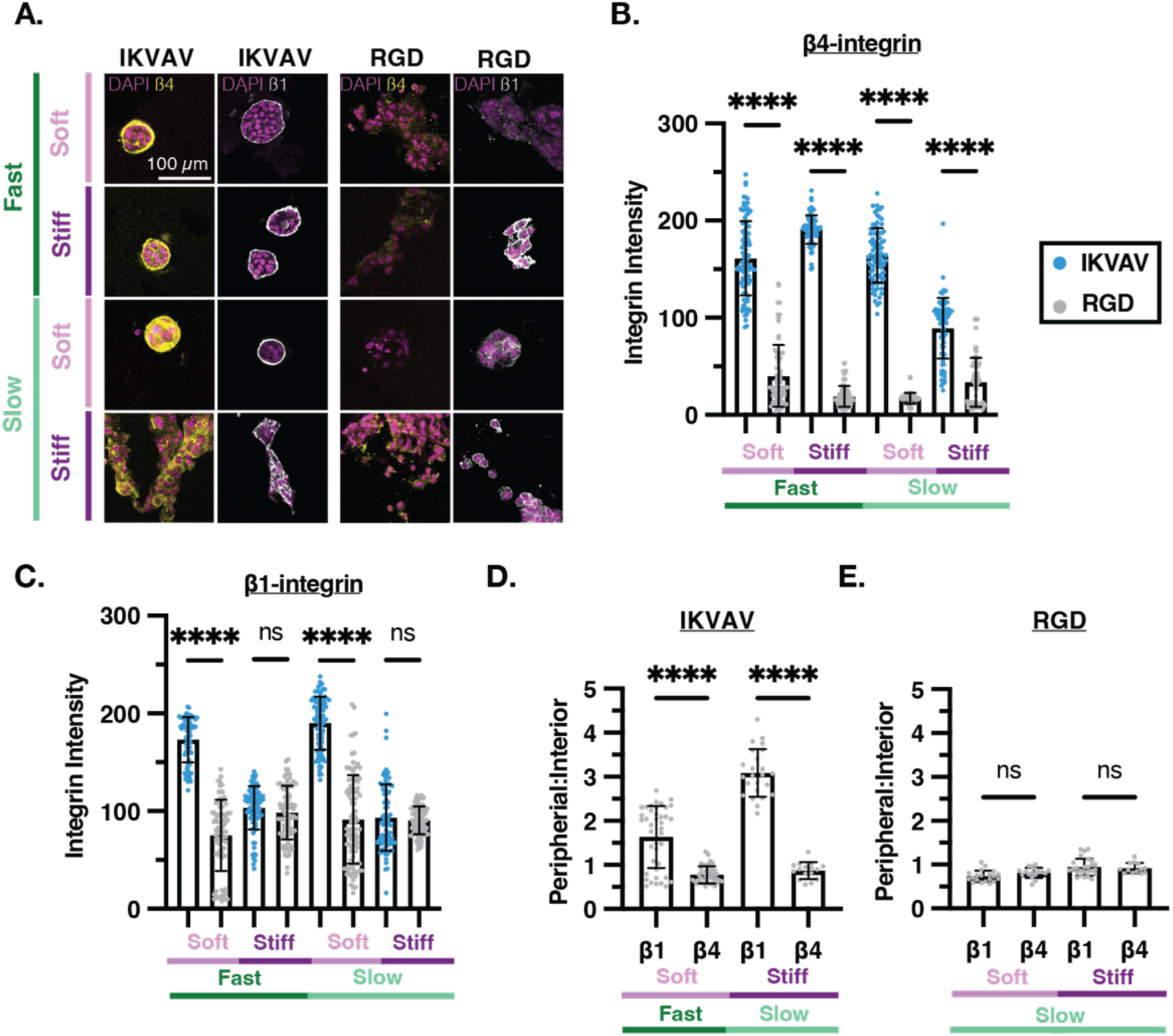
Polarization of β1 and β4 integrins associated with acinar development. (**A**) Representative confocal images of MCF10As encapsulated in high concentration IKVAV-modified eBMs (*Left)* and high concentration RGD-modified eBMs (*Right).* From left to right: DAPI/β4-integrin and DAPI/β1-integrin. (**B**) Quantification of β1-integrin intensity between IKVAV- and RGD-modified eBMs. (**C**) Quantification of β4-integrin intensity between IKVAV- and RGD-modified eBMs. (**D**) Quantification of peripheral to inner β1- and β4-integrin intensity within IKVAV-modified, fast-relaxing and soft, and slow-relaxing and stiff eBMs. (**E**) Quantification of peripheral to inner β1- and β4-integrin intensity within RGD-modified, fast-relaxing and soft, and slow-relaxing and stiff eBMs. All scale bars = 100 µm. Data are shown as mean ± SD of three biological replicates (n = 3, 20-50 images per replicate) unless otherwise indicated. Intensity measurements show all individual measurements. Peripheral: Interior measurements were taken from n > 30 images total over three biological replicates. Statistical significance was determined by one-way ANOVA and post hoc multiple comparison tests for integrin intensity and peripheral: interior ratio. * indicates p < 0.05, ** p < 0.01, *** p < 0.001, **** p < 0.0001, ns= not significant.

To determine the relative abundance of β1 and β4 integrins, we quantified the mean intensity from confocal microscopy images. We saw a significantly higher β1-integrin intensity in IKVAV-modified, fast-relaxing, soft and IKVAV-modified, slow-relaxing, soft eBMs compared to their RGD-modified counterparts (Fig. 5B). We did not observe a significant difference in β1-integrin intensity in IKVAV-modified, stiff eBMs compared to their RGD-modified counterparts (Fig. 5B). Conversely, β4-integrin intensity was significantly higher in all IKVAV-modified eBMs compared to their RGD eBM counterparts. Along with this, IKVAV-modified, slow-relaxing, stiff eBMs, which was the only group to induce malignant phenotypes and invasion, had the lowest β4-integrin intensity when compared to other IKVAV-modified eBMs (Fig. 5C, Fig. S3A).

To compare the extent of polarization of each integrin subunit, we quantified the relative localization of peripheral versus interior integrin intensity. Notably, there was greater peripheral localization of β1-integrin in comparison to β4-integrins within IKVAV-modified eBMs (Fig. 5D). In both IKVAV-modified, fast-relaxing, soft and IKVAV-modified, slow-relaxing, and stiff eBMs, β1-integrin was most abundant on the cluster periphery, while β4-integrin was found at both the periphery and the interior of the cluster (Fig. 5D). In contrast, there were no significant differences in the peripheral: interior localization of β1- and β4-integrins in RGD-modified, slow-relaxing eBMs (Fig. 5E). Based on strong localization of both β1- and β4-integrin to the basal side of acini in IKVAV-modified eBMs, we hypothesized that there is a balance in activity between β1- and β4-integrins in this context. To explore this further, we investigated the downstream signaling events associated with β1- and β4-integrin activation in IKVAV- and RGD-modified eBMs.

### Hemidesmosome Formation is Promoted in IKVAV-modified, Fast-Relaxing, Soft Matrices

Laminin 332-bound α6β4 integrins are known to cluster at the cell membrane, enabling MECs to form hemidesmosomes that provide stable attachment to the ECM (60). The adaptor protein plectin connects the cytoplasmic tail of clustered β4-integrins to the keratin intermediate filament network (Fig. 6A, left). Previous research has identified that hemidesmosome-dependent polarity, along with ECM composition, correlates with the growth rate and acinar formation of MECs (14,61).

**Fig. 6.**
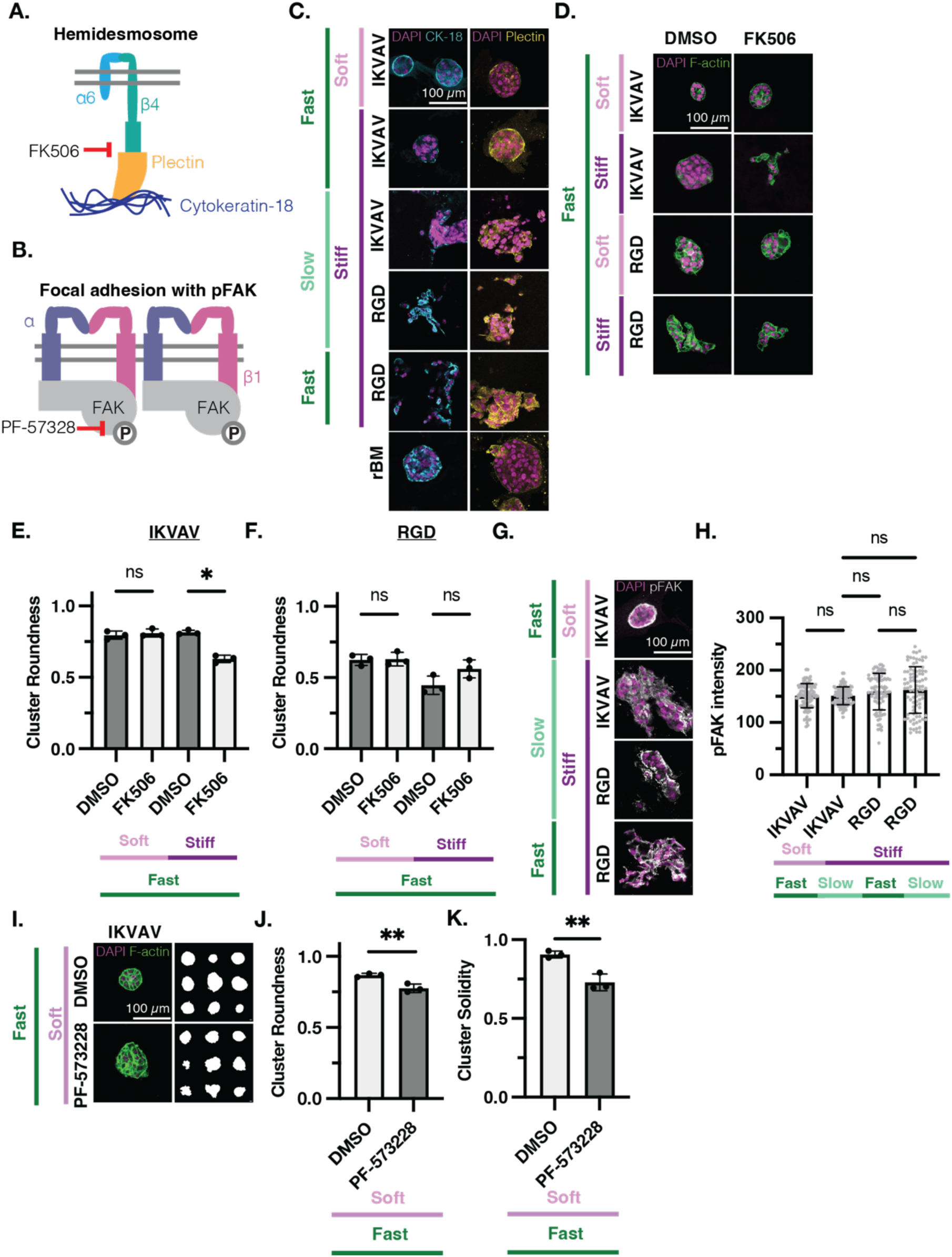
A Balance of β1 and β4 Integrin Signaling Leads to Acinar Formation in Modified eBMs. (**A**) Schematic of hemidesmosome structures and inhibition with FK506. (**B**) Schematic of focal adhesion and FAK phosphorylation and inhibition. (**C**) Representative confocal images of MCF10As encapsulated in IKVAV-modified fast-relaxing and soft, fast-relaxing and stiff, and slow-relaxing and stiff eBMs; RGD-modified slow-relaxing and stiff and fast-relaxing and stiff eBMs; rBM. From left to right: DAPI/Cytokeratin-18 and DAPI/Plectin. (**D**) DMSO (control) or FK506 (hemidesmosome inhibitor) treatment. Representative confocal images of MCF10As encapsulated in IKVAV-modified, fast-relaxing, soft and stiff eBMs; RGD-modified, fast-relaxing, soft and stiff eBMs. (**E**) Quantification of cluster roundness of IKVAV-modified eBMs from *D.* (**F**) Quantification of cluster roundness of RGD-modified eBMs from *D*. (**G**) Representative confocal images of MCF10As encapsulated in IKVAV-modified, fast-relaxing and soft, and slow-relaxing and stiff eBMs; RGD-modified, slow-relaxing and stiff and fast-relaxing and stiff eBMs. DAPI/pFAK. (**H**) Quantification of panel *G.* **I.** DMSO (control) or PF-573228 (FAKi) treatment. Representative confocal images of MCF10As encapsulated in IKVAV-modified, fast-relaxing and soft eBMs. From left to right: DAPI/F-actin and representative outlines. (**J**) Quantification of cluster roundness from panel *I.* (**K**) Quantification of cluster solidity from panel *I*. All scale bars = 100 µm. Data are shown as mean ± SD of three biological replicates (n=3, 20-50 images per replicate) unless otherwise indicated. Intensity measurements show all individual measurements. Statistical significance was tested by a Kruskal-Wallis test followed by Dunnett’s multiple testing correction for cluster roundness, by one-way ANOVA and post hoc multiple comparison tests for pFAK intensity, and a t-test for cluster solidity (p-value < 0.05). * indicates p < 0.05, ** p < 0.01, *** p < 0.001, **** p < 0.0001, ns= not significant.

We found that plectin and cytokeratin-18 localized to the basal side of acini in the IKVAV-modified, soft and stiff fast-relaxing eBMs, similar to the rBM controls (Fig. 6B). The presence of clustered β4-integrins, plectin and cytokeratin-18 indicate that hemidesmosome formation is likely a key contributor to the acinar phenotype in the IKVAV-modified, fast-relaxing eBMs. In IKVAV-modified, stiff, slow-relaxing eBMs and RGD-modified eBMs, which all yield malignant phenotypes, we did not see basal localization of cytokeratin-18 and plectin. Instead, there was diffuse distribution of these proteins throughout the cell clusters (Fig. 6B).

To investigate the role of hemidesmosomes in maintaining the acinar phenotype, we inhibited their formation in MCF10A cells cultured in modified eBMs. FK506, a calcineurin inhibitor, disrupts hemidesmosomes by phosphorylating the β4-integrin cytoplasmic tail, (Fig. 6A, right) (62), freeing α6β4 to activate pro-invasive PI3K, Akt, and MAPK signaling (63). In IKVAV-modified, fast-relaxing, soft eBMs, FK506 treatment had no significant effect on cluster morphology (Fig. 6C, D), suggesting that a IKVAV-modified, fast-relaxing, soft matrix does not promote invasion regardless of hemidesmosome status. However, in stiff, fast-relaxing IKVAV-modified eBMs, inhibition of hemidesmosomes led to a marked decrease in roundness and the emergence of an invasive phenotype (Fig. 6C, D), indicating that hemidesmosomes play a critical role in restricting invasion even in a stiff environment. Notably, RGD-modified eBMs showed no significant changes in phenotype or cluster roundness upon FK506 treatment (Fig. 6C, E), demonstrating that this effect is specific to β4-integrin interactions in the IKVAV-modified matrix. Together, these findings suggest that hemidesmosomes help preserve the acinar phenotype and prevent invasion despite elevated matrix stiffness, while their loss enables invasive behavior.

### Phosphorylated FAK Contributes to Acinar Morphogenesis in IKVAV-modified eBMs

To examine β1-integrin signaling, we assessed phosphorylated focal adhesion kinase (FAK) (pY397) in modified eBMs. β1-integrin activation leads to clustering at the membrane (64,65), recruitment and phosphorylation of FAK at Y397 (66–68), and ultimately regulation of MEC responses to physical cues (69,70) (Fig. 6F). Surprisingly, we observed strong pFAK intensity across all eBM conditions, regardless of stiffness or stress relaxation (Fig. 6G, H), with RGD-modified eBMs having more pFAK intensity overall compared to IKVAV-modified eBMs. This finding was unexpected, as phosphorylation of FAK has predominantly been associated with stiff matrices (17, 69, 71). FAK inhibition led to a significant decrease in cluster roundness in IKVAV-modified, fast-relaxing, soft eBMs, indicative of a disruption in acinus formation (Fig. 6I, J, K). We also found that FAK inhibition caused significant changes in the surface roughness of the cell clusters, quantified by solidity (Fig. 6J). These findings reveal a role for FAK signaling in acinar morphogenesis, demonstrating that FAK signaling is crucial not only in stiffer matrices but also in softer environments.

### IKVAV eBMs Promote Significant Endogenous Laminin Deposition Throughout Culture Period

Inspired by reports that cells in engineered hydrogels can deposit matrix proteins dependent on the initial crosslinking density of the hydrogel (72), we investigated whether MCF10A clusters in modified eBMs secrete basement membrane proteins, potentially aiding acinus formation under specific conditions. As laminin is the most abundant protein of the basement membrane, we stained clusters in the modified eBMs with pan-laminin and laminin 332 antibodies. Since IKVAV is derived from laminin 111 (47,51,73) laminin 332 should reflect only the laminin produced endogenously by cells.

Strikingly, laminin 332 deposition was significantly increased in IKVAV-modified eBMs compared to RGD-modified eBMs (Fig. 7A, B). In IKVAV eBMs, laminin was predominantly localized adjacent to the basal side of the cell clusters (Fig. 7A), with IKVAV-modified, fast-relaxing, soft eBMs exhibiting significantly higher laminin deposition than their slow-relaxing, stiff counterparts (Fig. 7B). In contrast, laminin deposition in RGD-modified eBMs was sparse and primarily confined within the cell clusters rather than surrounding them (Fig. 7A), and the overall levels were markedly lower than those observed in IKVAV-modified eBMs (Fig. 7B). We observed similar trends when staining with a pan-laminin antibody (Fig. S4A,B).

**Fig. 7.**
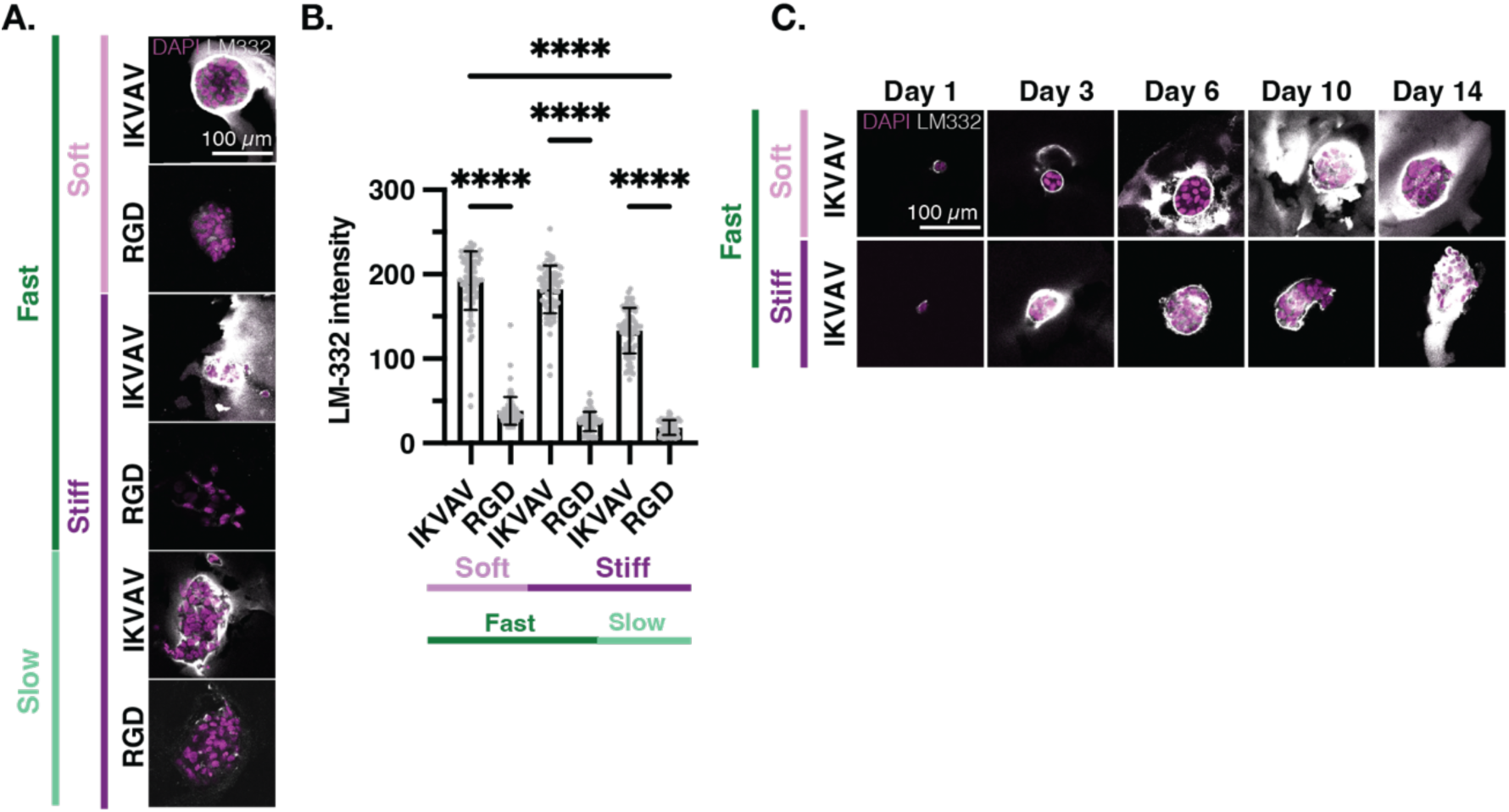
Only IKVAV eBMs Support Significant Endogenous LM-332 Deposition. (**A**) Representative confocal images of MCF10As encapsulated in IKVAV- and eBMs. DAPI/LM-332. (**B**) Quantification of LM-332 intensity from Panel *A.* (**C**) Representative confocal images of MCF10As encapsulated in IKVAV-modified fast-relaxing and soft, and fast-relaxing and stiff eBMs. Images taken at Day 1, 3, 6, 10 and 14. DAPI/LM-332. All scale bars = 100 µm. Data are shown as mean ± SD of three biological replicates (n=3, 20-50 images per replicate) unless otherwise indicated. Intensity measurements show all individual measurements. Statistical significance was tested by one-way ANOVA and post hoc multiple comparison tests for LM-332 intensity. * indicates p < 0.05, ** p < 0.01, *** p < 0.001, **** p < 0.0001, ns= not significant.

Next, we investigated the timescale of laminin deposition by immunostaining and imaging IKVAV-modified eBMs at intermediate time points within the culture period.

Laminin was deposited as early as Day 1, and was continuously produced throughout the 14-day experimental time course (Fig. 7C). Laminin 332 deposition between IKVAV-modified, fast-relaxing soft and stiff eBMs was similar by the end of the time points (Fig. 7C). The strong laminin 332 deposition in IKVAV eBMs, but not in RGD-modified eBMs, suggests that MCF10A clusters in modified eBMs not only respond to their environment but also contribute to endogenous laminin 332 deposition, potentially enhancing cellular polarization and functionality over time, depending on the condition. Furthermore, this outcome underscores the critical role of the initial eBM condition provided to the cell, highlighting its influence on both the quantity and the nature of the endogenous proteins synthesized.

Collectively, these results indicate that the IKVAV motif alone is sufficient to promote mammary acinar morphogenesis, provided the matrix exhibits an appropriate combination of mechanical properties. Furthermore, a balanced engagement of β1- and β4-integrins appears to be crucial for successful acinar development in IKVAV-modified eBMs. However, this process is contingent upon the matrix being sufficiently dynamic to support both laminin deposition and integrin clustering.

## Discussion

We have designed a tunable 3D matrix system with definable parameters for culturing MECs that enables mammary acinar morphogenesis without rBM. Importantly, this system allows for the independent manipulation of both biochemical and mechanical properties, effectively decoupling their effects, which has been a challenge with traditional rBM systems. IKVAV-modified, soft, fast-relaxing eBMs, designed to mimic rBM, promoted polarized acini in MECs. We also observed an acinar-like phenotype in both the IKVAV-modified, fast-relaxing, stiff eBMs and IKVAV-modified, slow-relaxing, soft eBMs. This novel platform overcomes limitations of existing synthetic systems, which have either required exogenous matrix components or lacked the independent control over matrix mechanics necessary for complete acinar polarization. Self-assembling peptide hydrogels (SAPHs) like PeptiGel® have been explored for rBM-free mammary epithelial cell culture, but achieving complete acinar maturation has required exogenous laminin supplementation (74). Additionally, while PEG hydrogels have shown promise for mammary acini formation through stiffness optimization and RGD peptide conjugation, they still lack independently tunable viscoelasticity (75). Moreover, our work demonstrates the substantial positive impact of IKVAV adhesion motifs on acinar morphogenesis in comparison to RGD motifs. We have also successfully cultured patient-derived tumor organoids in IKVAV-modified eBMs, demonstrating that modified eBMs can be versatile and applicable to other biological systems.

Prior reports have elegantly shown in rBM-based systems that matrix stiffness in the range of malignant breast tumors (1-5 kPa) drives invasion in MECs (14,15,71).

However, in our IKVAV-modified eBMs, tumor-like stiffness only induced invasion when the matrix was both slow-relaxing and stiff. Several factors may explain this difference. Firstly, the stress relaxation rates of the eBM play a crucial role in integrin clustering.

Fast-relaxing matrices facilitate integrin clustering more effectively compared to slow-relaxing matrices (77,78). Secondly, we are independently modulating the stress relaxation properties of the eBMs, a mechanical perturbation that has not been previously characterized in clusters grown from single MECs in rBM-free systems.

Finally, while the use of a rBM-free system allows us to unravel the contributions of different ECM components in a reductionist manner, there will be inherent differences from experiments performed with the complexity of rBM components. These could include the presence of growth factors and other signaling molecules from rBM products, which have been reported to affect cell culture studies as well as contributions from the complex interactions of basement membrane structural proteins that are not recapitulated by peptide-modified alginate networks (31–33).

An invasive phenotype was present in RGD eBMs regardless of their mechanical properties. The difference in phenotypes between the IKVAV-modified and RGD-modified eBMs clearly demonstrates the substantial impact of the chosen peptide motif on acinar development. Different integrin subunit combinations bind the IKVAV and RGD motifs, and thus induce different integrins to cluster, subsequently forming distinct adhesion complexes, leading to the emergence of the observed phenotypes. In MECs, β1- and β4-integrins play a role in both polarized and invasive phenotypes, with the ECM modulating their functions (79). For instance, β4-integrins promote invasion via the PI3K pathway (80) but are also involved in hemidesmosome formation and acinar development. The role of β1-integrins is more complex; blocking β1 can reverse malignant phenotypes (15), while α2β1 integrins activate Rac and promote epithelial polarization (81). Notably, α2β1 is enriched in healthy mammary tissue but downregulated in tumors (82,83).

We found that β4-integrin consistently localized to the basal side of acinar-like clusters. However, in invasive clusters, like in the IKVAV-modified, slow-relaxing and stiff eBMs, β4-integrin was randomly dispersed. We anticipate that differences in β4-integrin clustering explain the phenotypic differences we observed. Cells cluster integrins more effectively in fast-relaxing matrices (38) due to physical remodeling of the alginate network, thus facilitating integrin clustering and hemidesmosome formation.

The lack of acinar morphogenesis in alginate gels with low IKVAV concentration further supports this hypothesis, since too few integrin-ligand complexes exist to effectively cluster for downstream signaling. Interestingly, inhibition of both FAK and hemidesmosome formation disrupted acinar development in IKVAV-modified eBMs.

Since FAK is linked to β1-integrin signaling and hemidesmosomes to β4-integrin signaling, these results demonstrated that both β1- and β4-integrin signaling is critical for the acinar phenotype in these IKVAV-modified eBMs. Specifically, β4-integrin’s role in hemidesmosome formation is required for acinar development in IKVAV-modified, fast-relaxing, stiff eBMs. Prior work has shown that enhanced stiffness can drive the invasive phenotype via PI3K and Rac1 signaling from the tail of β4 integrin if matrix mechanical conditions prevent hemidesmosome formation (14,59,84,85). Our results similarly show that matrices that hinder β4-integrin clustering, such as stiff and slow-relaxing matrices or those with low IKVAV concentrations, promote the formation of invasive clusters. Fast stress relaxation, even in stiff matrices, enables hemidesmosome formation and prevents invasion. Additionally, phosphorylated FAK staining, downstream of β1-integrin activity, revealed strong signal intensity in the acini formed in IKVAV-modified, fast-relaxing, soft eBMs. Inhibition of FAK signaling resulted in a significant reduction in cluster roundness and a disruption of cluster organization. Our findings suggest that the acinar phenotype in IKVAV eBMs arises from a combination of mechanical and cell adhesion factors that promote a balance of β1- and β4-integrin activity.

This study observed increased endogenous laminin deposition in IKVAV-modified, fast-relaxing, soft eBMs compared to their slow-relaxing, stiff counterparts highlights the role of matrix compliance in facilitating ECM protein deposition. A more compliant, fast-relaxing matrix may alleviate physical constraints, thereby enhancing laminin deposition, consistent with previous findings (72,86). Furthermore, the observed laminin deposition in both fast-relaxing stiff and soft IKVAV-modified eBMs, contrasted with its absence in RGD-modified eBMs, underscores the critical influence of the adhesion peptide motif on matrix deposition and cellular phenotype. Elevated laminin levels are likely to drive acinar formation through multiple mechanisms, including promotion of hemidesmosome formation through enhanced integrin clustering and protection of the nucleus from mechanical deformation (14,87). Consistent with this, hemidesmosomes were evident in IKVAV-modified eBMs, while inhibition of their formation resulted in an invasive phenotype exclusively in fast-relaxing, stiff IKVAV-modified eBMs. Notably, no phenotypic differences were observed in the RGD-modified eBM control or treated groups, further highlighting the unique role of IKVAV in promoting acinar-like organization. In addition to hemidesmosome formation, laminin deposition may contribute to nuclear stability. Recent studies suggest that high laminin levels form a keratin-laminin shield, protecting the nucleus from mechanical deformation and dampening cellular responses to mechanical cues (87). This protective mechanism may explain how increased laminin deposition in fast-relaxing, stiff IKVAV-modified eBMs supports nuclear integrity, ultimately favoring acinar morphology. Finally, a cascading chain of events leads to α6β4 integrin clustering: α2β1 integrin activation drives laminin-332 deposition and organization, which in turn creates additional binding sites, culminating in α6β4 integrin clustering (61,81,88). This integrin-mediated positive feedback loop likely plays a pivotal role in reinforcing acinar phenotypes in IKVAV-modified eBMs (79). Together, these findings emphasize the interplay between matrix mechanics, laminin deposition, and integrin signaling in driving acinar formation.

While this study focused on IKVAV and RGD eBMs, additional peptides, such as adhesion motifs derived from collagen IV, could be conjugated to an alginate matrix to better mimic the components of rBM, given that collagen IV constitutes 30% of the basement membrane (32). This approach would refine our understanding of the specific peptides required for mammary morphogenesis. Furthermore, combining multiple modified alginates would present various peptides to cells, enabling more complex studies of ECM ligand interactions and their effects on mammary epithelial cell phenotype.

Although our reductionist matrix allows precise control over individual mechanical and biochemical cues—offering a powerful tool to dissect their distinct roles—it does not replicate the full complexity of rBM, which contains over 2,000 proteins that may contribute synergistically to certain biological processes (9,32). While this complexity poses challenges for reproducibility and mechanistic studies, it also presents utility in applications where an undefined yet functionally rich microenvironment is beneficial.

Our approach, therefore, complements existing models by enabling controlled investigations into ECM properties that would be difficult to isolate in traditional rBM-based systems. Additionally, this platform has the potential to be extended to other cell types requiring precise modulation of multiple mechanical and biochemical cues to understand their individual contributions.

Beyond mammary epithelial morphogenesis, our eBM system offers broad utility for studying mechanotransduction, ECM cues, and integrin signaling across diverse biological contexts. The modularity of this platform enables the incorporation of additional ECM-derived motifs to further refine basement membrane mimicry, paving the way for more physiologically relevant in vitro models. Moreover, the successful culture of patient-derived tumor organoids in our system underscores its potential for translational applications, including precision medicine and therapeutic screening. By bridging the gap between reductionist synthetic matrices and the complexity of native basement membranes, this platform provides a powerful tool to unravel fundamental mechanisms governing tissue organization, disease progression, and cellular responses to the microenvironment.

## Methods

### Materials

Purified Pronova UPVLVG and Pronova LF20/40 alginates were purchased from Novamatrix. RGD, YIGSR, and IKVAV peptides were obtained from Peptide 2.0. Full Peptide sequences and their molecular weight are as follows: GGGGRGDSP (758.75 g/mol), CQAASIKVAV (989.19 g/mol), CDPGYIGSR (967.06 g/mol). Growth factor reduced Matrigel™ was purchased from Corning (Cat. #354230).

### Alginate Preparation, Conjugation and Characterization

Pronova LF20/40 alginate was purified by dialysis (3500 MWCO tubing) against Milli-Q water for four days, frozen and lyophilized until dried. LF20/40 (high molecular weight) or VLVG (low molecular weight) alginate was dissolved in 0.1 MES buffer overnight. Carbodiimide chemical reactions were carried out in a 0.1 M MES buffer (pH = 6.5) solution in order to modify alginate chains with their respective peptides. Peptides (either RGD, YIGSR, or IKVAV) were conjugated to the alginate backbone via their terminal amine. Separate batches of alginate were made for each respective peptide (RGD, YIGSR, or IKVAV). 1-ethyl-(dimethylaminopropyl) carbodiimide (EDC) was used to form amide linkages between amine containing-molecules of the peptides and the carboxylate moieties on the alginate polymer backbone. N-hydroxy-sulfosuccinimide (sulfo-NHS) was utilized as a co-reactant to stabilize the reactive EDC intermediates and enhance conjugation efficiency. Sulfo-NHS, EDC, and respective peptides were added quickly and sequentially to an alginate solution in order to achieve homogeneously modified alginate. The reaction was carried out for 20 hours, then quenched with 100X hydroxylamine (125 mg/g alginate). Activated charcoal was added to each batch of alginate and mixed for 30 minutes, allowing the charcoal to settle afterward. The alginate was then filtered twice, first through a 0.8 µM filter and then through a 0.2 µM filter. After filtration, the alginate was frozen at -80°C before lyophilization. The samples were lyophilized at -50°C for 72 hours. Once complete, the alginate was reconstituted under sterile conditions in DMEM/F12 basal media and used at a final concentration of 1% in the gels. The degree of peptide conjugation was approximated via ^1^H-NMR spectroscopy (500 MHz, Bruker, Germany). The ^1^H-NMR spectra and the corresponding peak assignments for each sample are provided in the supplementary materials (Fig. S1, Table S1).

### Rheology

Rheology experiments were conducted using a stress-controlled Anton Paar MCR 502 rheometer. Briefly, the gels were fabricated and deposited directly onto the bottom plate, and a 20mm plate was then lowered slowly to contact the gel. Mineral oil was deposited to the periphery of the gel to avoid gel dehydration. A time sweep was conducted (1 Hz, 1% strain, 37 °C) and the storage (G’) and loss modulus (G”) were recorded overtime. From this data, the complex modulus (G*) was calculated (Equation 1), and subsequently the elastic modulus (E) was calculated (Equation 2). A Poisson’s ratio (*v*) was assumed to be 0.5. Once the storage modulus reached an equilibrium value, a stress-relaxation test was subsequently performed at 10% strain. The stress-relaxation time was defined as the time taken for the maximum stress to relax to half of its initial value.

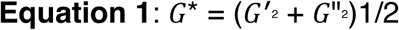

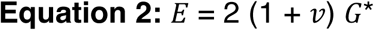

### Mammary Epithelial Cell Culture

Immortalized, non-tumorigenic MCF10A mammary epithelial cells were obtained from ATCC and cultured via established protocols (1).

Cells were cultured as a monolayer in 2D tissue culture flasks until time for encapsulation. The following growth media recipe was used: DMEM/F12 (ThermoFisher Scientific) basal media was supplemented with 5% horse serum (ThermoFisher Scientific), 1% penicillin/streptomycin (Thermo Fisher), 20 ng/mL EGF (Peprotech), 0.5 mg/mL hydrocortisone (Sigma), 100 ng/mL cholera toxin (Sigma), and 10 µg/mL insulin (Sigma). Prior to encapsulation, MCF10As cells were washed with PBS, trypsinized (0.05% trypsin/EDTA, ThermoFisher Scientific), spun down (2,000 rcf, 5 minutes), counted and resuspended in DMEM/F12 to create a single-cell suspension with a final concentration in the hydrogels of 50,000 cells/mL.

### eBM Formation

Alginate (1% w/v final concentration) in DMEM/F12 and a single-cell suspension (final concentration of 50,000 cells/mL) in growth media were mixed in a Luer lock syringe. Calcium sulfate was used to crosslink alginate matrices. A calcium sulfate slurry was mixed with DMEM/F12 and added to a second Luer lock syringe. Final calcium concentrations for soft and stiff matrices were 4 mM and 22 mM for LF20/40 matrices, and 7 mM and 25 mM for VLVG matrices, respectively. The two syringes were connected via a coupler and the solutions were mixed several times. The hydrogel solution was then quickly deposited into well plates of a 48-well plate (∼500 μL per well). Hydrogels were incubated at 37°C for two hours before adding growth media, and were then cultured for two weeks. Media was changed every 1-2 days.

### MCF10A Cell Encapsulation in rBM

For MCF10A encapsulation into rBM, Matrigel (Corning, Cat. #354230) was mixed with a single-cell suspension (DMEM/F12) and gels were deposited into the wells of a 48-well plate (∼500 μL per well). rBM matrices were incubated at 37°C for two hours before adding growth media, and were then cultured for two weeks. Media was changed every 1-2 days.

### PDM-350 Culture

HCM-CSHL-0261-C50 tumor organoids were obtained from ATCC and cared for via established ATCC protocols. Advanced DMEM/F12 was supplemented with Organoid Growth Kit 1F (ACS-7105), L-Glutamine (ATCC, 30-2214), HEPES (Thermo Fisher, 15630080), B-27 Supplement (Thermo Fisher, 17504-044), and HA-R-Spondin1-Fc 293T (RSPO1) conditioned media (Trevigen, Cat# 3710-001-01) for culture media. Organoids were thawed, centrifuged into a pellet, and resuspended in 100-200 μL Matrigel for culture. 10 μL droplets or domes were dispensed in a 6-well plate with about 10 droplets per well. The 6-well plate was inverted and placed in the 37 ℃ incubator for 30 minutes. After incubation, 6-well plates were placed upright and 2.0 mL growth media was added per well. Media was changed every 1-2 days.

### PDM-350 Encapsulation in eBMs and rBM

Organoids were scraped from 6-well plate, centrifuged into a pellet, and resuspended in TrypLE (Thermo Fisher, #12604013) to generate single cells. The single cells were quenched, pelleted again, and resuspended in growth media or Matrigel for encapsulation.

### Inhibitor Studies

All inhibitors were diluted in growth media and added 1-day post-encapsulation. DMSO was used as a vehicle control for all inhibitors. FK506 (Cayman Chemical, 10007965) was used at a working concentration of 1 μM. PF-573228 (FAK inhibitor, Cayman Chemical) was used at a working concentration of 5 μM.

### Immunofluorescence Staining and Confocal Microscopy

Hydrogel samples were fixed in 4% paraformaldehyde in PBS for 45 minutes at 37°C. Samples were then washed twice with Dulbecco’s phosphate-buffered saline (DPBS) with Ca^2+^/Mg^2+^ for 15 min, followed by an overnight incubation in 30% v/v sucrose-cPBS solution to dehydrate the hydrogels. The hydrogels were then placed in a solution of equal volumes OCT (Tissue-Tek) and 30% sucrose-cPBS for four to six hours. After incubation, the solution was removed and the gels were placed individually into cryomolds with OCT and then frozen at -20°C. The samples were then sectioned with a cryostat (Lecia CM1950, 40 µm) for immunostaining. Sections were washed with PBS, then incubated in blocking buffer (10% goat serum (ThermoFisher Scientific), 1% bovine serum albumin (Sigma), 0.1% Triton X-100 (Sigma), and 0.3 M glycine (Sigma)) for 1 hour before addition of primary antibodies. Samples were incubated with primary antibodies overnight, then rinsed with blocking buffer twice. Alexa Fluor 488–phalloidin *1:100 dilution* (Thermo Fisher, A12379), 4′,6-diamidino-2phenylindole (DAPI; 1 μg ml−1), and fluorescently conjugated secondary antibodies were diluted in the blocking buffer and added to slides for 1 hour at RT. *Specific primary antibodies used:* GM130 *1:100* (BD biosciences, 61083), β4-integrin *1:150* (Life Sci. Tech., MA5-17104), β1-integrin *1:200* (Thermo, 14-0299-82), Plectin *1:100* (Abcam, EPR26993-49), Keratin-14 *1:234* (Abcam, AB312312-1001),

Laminin 332 *1:100* (P3H9, DSHB). *Specific secondary antibodies used*: Alexa Fluor IgG1 goat anti-mouse, 488 (Thermo Fisher, A21121), Alexa Fluor IgG1 goat anti-mouse 555 (Thermo Fisher, A21127), and Alexa Fluor IgG2 goat anti-rabbit, 488 (Thermo Fisher, A11008). Slides were imaged with a 25x objective on a Leica SP8 confocal laser scanning microscope.

### Image Analysis

ImageJ (NIH) was utilized for image analysis to evaluate the roundness, area, and invasion of cell clusters, along with the localization of specific markers. At least three replicates (20-50 images per replicate) were used per condition unless otherwise stated in figure caption. Confocal images were thresholded and masked using ImageJ software to create outlines and analyze shape metrics of each cluster. Using this method, roundness, or the inverse of the aspect ratio (4*area/(π*major_axis^2) and area (µm^2^) were calculated. The average of each experimental replicate is displayed as one datapoint. Fluorescence intensity analysis involved calculating the mean intensity per unit area (µm^2^) for each cluster to facilitate comparisons across conditions. Interior-to-peripheral intensity measurements were performed by defining a 10-micron boundary surrounding the cell cluster to represent the “basal membrane” for peripheral intensity measurements. Interior intensity measurements were taken from the region within this boundary, excluding a 5-micron margin to avoid overlap with the peripheral zone. The percentage of invasive clusters was determined by classifying a cluster as invasive if it displayed a distinct protrusion, characterized by a discontinuous change in curvature at the resolution of the brightfield image (approximately 1 μm).

### Statistical Analysis

All data was analyzed in GraphPad Prism v9.3.1 software using the tests described in the respective figure captions. P-values of less than 0.05 were considered statistically significant. Unless otherwise noted, graphs present three biological replicates per condition (n=3), with each replicate representing the average at least 20-50 images.

## Acknowledgements

The P3H9 antibody, developed by Fred Hutchinson Cancer Research Center (Wayner EA), was obtained from the Developmental Studies Hybridoma Bank, created by the NICHD of the NIH and maintained at The University of Iowa, Department of Biology, Iowa City, IA 52242. We acknowledge the use of the NRI-MCDB Microscopy Facility at the University of California, Santa Barbara. Dr. Jerry Hu is gratefully acknowledged for running NMR and peak assignments.

## Author Contributions

Conceptualization: JAB, RSS

Methodology: JAB, RSS

Investigation: JAB, MDL, SMJ, AS, DIW, RSS

Visualization: JAB, AS, RSS

Analysis: JAB, MDL, SMJ, AS, RSS

Supervision: JAB, RSS

Writing—original draft: JAB, RSS

Writing—review & editing: JAB, SMJ, AS, DIW, RSS

## Data and materials availability

All data needed to evaluate the conclusions in the paper are present in the paper and/or the Supplementary Materials.

---

**Fig. S1.**
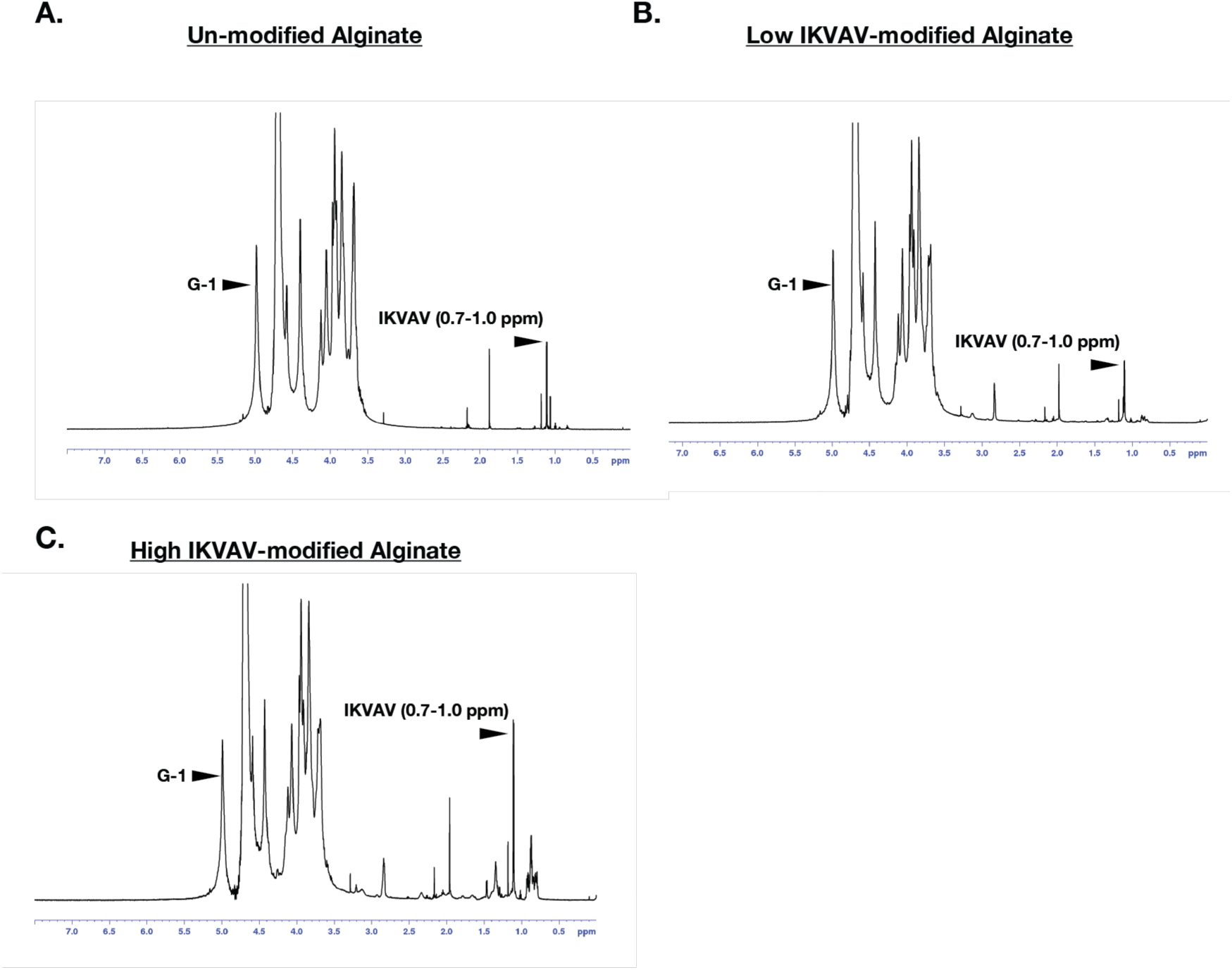
1H-NMR spectra of alginate and IKVAV-modified fast-relaxing alginate at varying concentrations. (A) Unmodified alginate. (B) Low concentration IKVAV-modified fast-relaxing alginate. (D) High-concentration IKVAV-modified fast-relaxing alginate. One sample per condition was analyzed. “G-1” denotes alginate peak analyzed and “IKVAV” denotes peptide peak(s) analyzed.

**Table S1.**
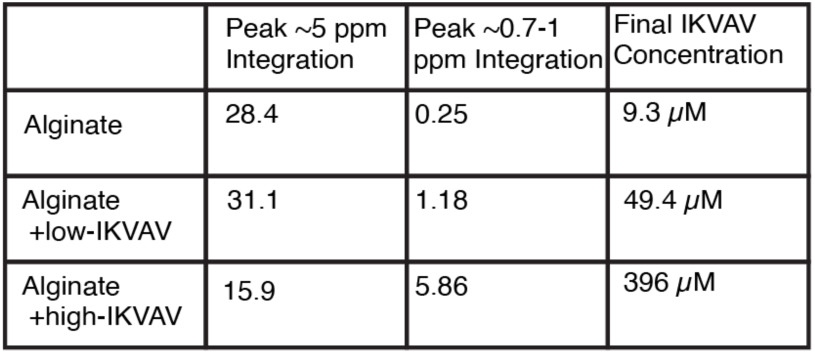
The integration of assigned 1H signals on IKVAV-modified alginate. The integration of assigned 1H signals on alginate (G-blocks, ∼5.1 ppm) and IKVAV peptide (0.7–1.0 ppm) signals are shown, enabling the calculation of molar peptide concentration relative to the alginate monomer count (Alg-CQAASIKVAV: 3 protons from Ile 3-Hγ, 3 protons from Ile 3-Hδ, and 12 protons from Val 2-Hγ, totaling 18 protons). The molar peptide concentration was calculated using the equation: M_p_=[71*M_a_*I_1_]/[18*I_5_]. M_a_ = 2.67 x 10^-4^ when using a 2% solution of alginate. I_1_ and I_5_ correspond to peaks at ∼0.7-1.0 ppm and ∼5.0 ppm, respectively.

**Fig. S2.**
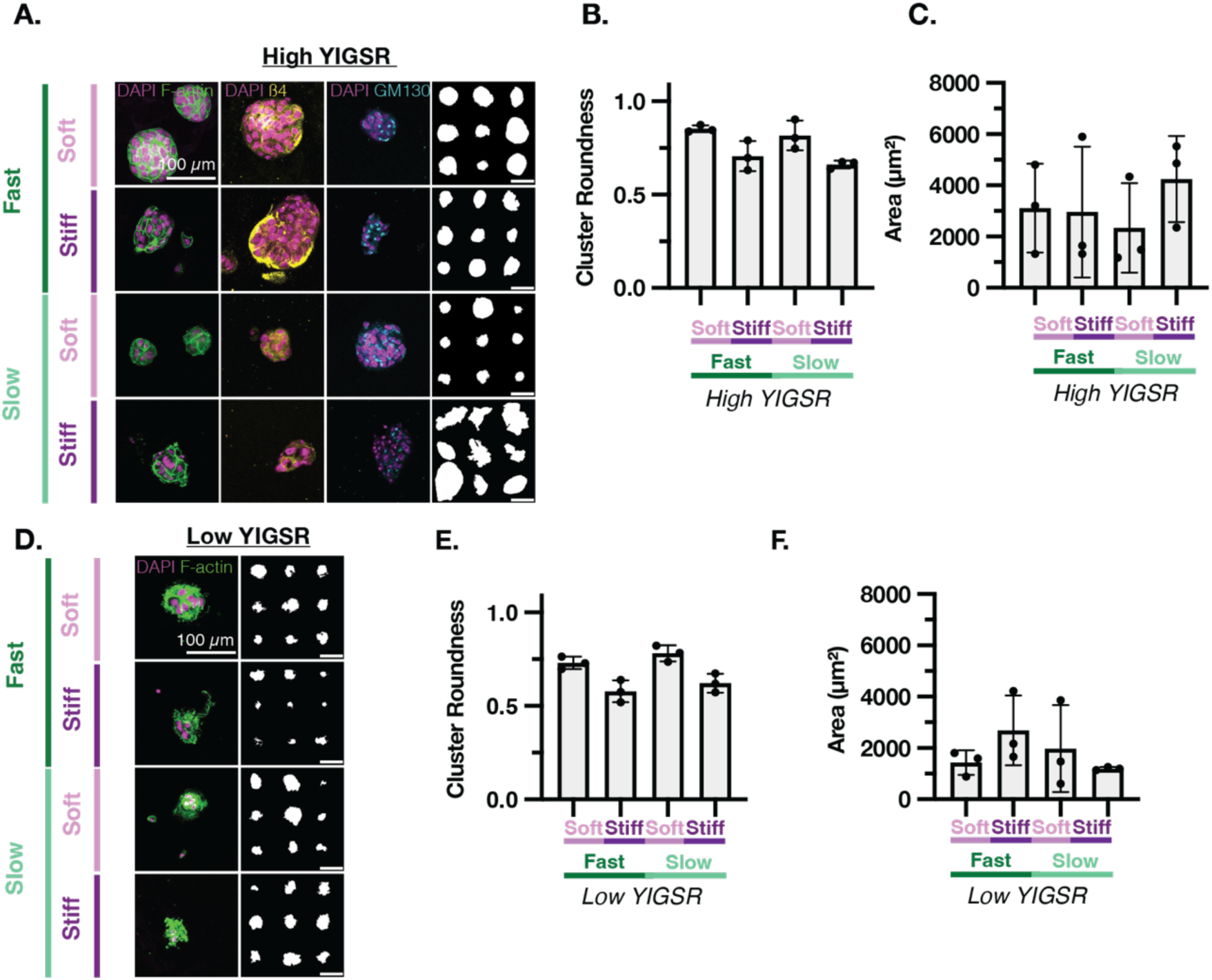
YIGSR eBMs Do Not Support Robust Mammary Morphogenesis. (A) Representative confocal images of MCF10As encapsulated in high concentration YIGSR-modified eBMs. From left to right: DAPI/F-actin, DAPI/β4-integrin, DAPI/GM130, and representative outlines of the respective condition. (B) Quantification of cluster roundness for high concentration YIGSR-modified eBMs. (C) Quantification of cluster area for YIGSR-modified eBMs. (D) Representative confocal images of MCF10As encapsulated in low concentration YIGSR-modified eBMs. DAPI/F-actin. Scale bars = *100* µm. (E) Quantification of cluster roundness for low concentration YIGSR-modified eBMs. All scale bars = 100 *µm*. Data are shown as mean ± SD of three biological replicates (n=3, 20-50 images per replicate) unless otherwise indicated. Statistical significance was tested by a Kruskal-Wallis test followed by Dunnett’s multiple testing correction for cluster roundness, and by one-way ANOVA and post hoc multiple comparison tests for cluster area. If no statistical significance indicator bars are shown, there were no significant differences (p-value > 0.05). * indicates p < 0.05, ** p < 0.01, *** p < 0.001, **** p < 0.0001.

**Fig. S3.**
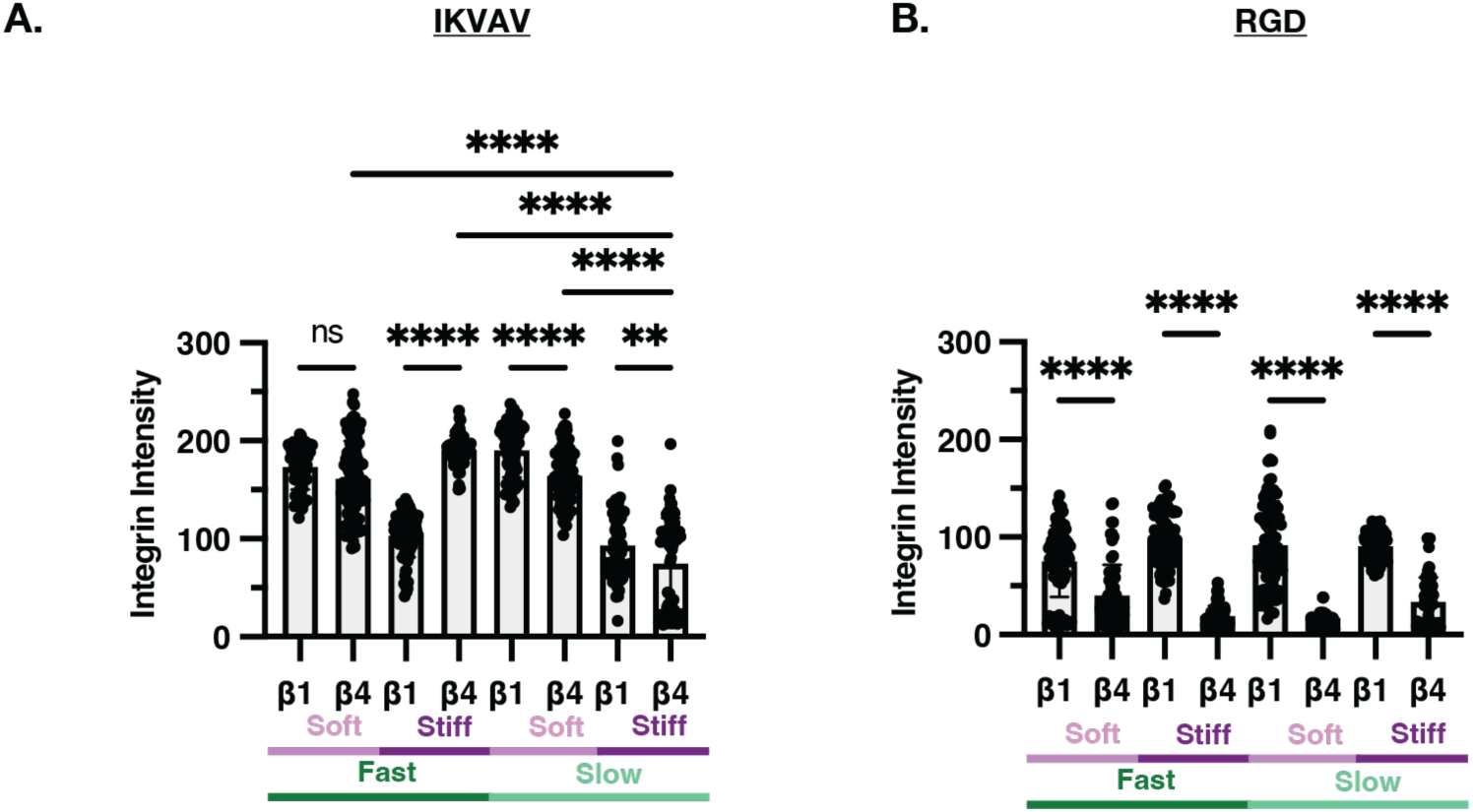
β1- and β4-Integrin expression is more balanced in IKVAV-modified eBMs as opposed to RGD eBMs. (A) β1 and β4 Integrin in IKVAV-modified eBMs. (B) β1 and β4 Integrin in RGD-modified eBMs. Data are shown as mean ± SD of three biological replicates (n=3, 20-50 images per replicate) unless otherwise indicated. Statistical significance was tested by one-way ANOVA and post hoc multiple comparison tests for integrin intensity (p-value < 0.05). * indicates p < 0.05, ** p < 0.01, *** p < 0.001, **** p < 0.0001, ns = not significant.

**Fig. S4.**
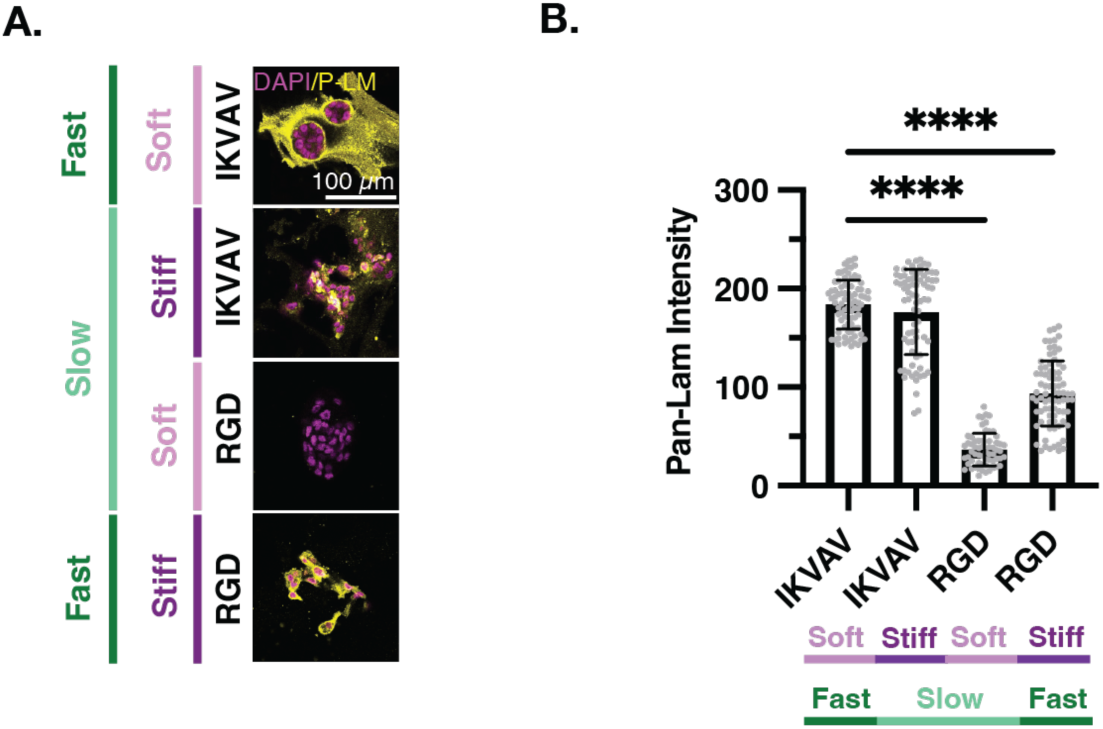
Pan-laminin intensity is significantly higher in IKVAV-modified eBMs. (A) Representative confocal images (day 14) of MCF10As encapsulated in IKVAV- and RGD-modified eBMs. DAPI/Pan-laminin. (B) Quantification of pan-laminin intensity from Panel *A.* All scale bars = 100 µm. Data are shown as mean ± SD of three biological replicates (n=3, 20-50 images per replicate) unless otherwise indicated. Intensity measurements show all individual measurements. Statistical significance was tested by one-way ANOVA and post hoc multiple comparison tests for pan-laminin intensity. * indicates p < 0.05, ** p < 0.01, *** p < 0.001, **** p < 0.0001, ns = not significant.

